# Cerebral atherosclerosis contributes to Alzheimer’s dementia independently of its hallmark amyloid and tau pathologies

**DOI:** 10.1101/793349

**Authors:** Aliza P. Wingo, Wen Fan, Duc M. Duong, Ekaterina S. Gerasimov, Eric B. Dammer, Bartholomew White, Madhav Thambisetty, Juan C. Troncoso, Julie A. Schneider, James J. Lah, David A. Bennett, Nicholas T. Seyfried, Allan I. Levey, Thomas S. Wingo

**Author notes:** Corresponding authors (T.S.W.), (A.I.L.), (N.T.S.), (A.P.W.), Emory University Center for Neurodegenerative Disease, 505K Whitehead Building, 615 Michael Street NE, Atlanta, GA 30322-1047.

## Abstract

Cerebral atherosclerosis is a leading cause of stroke and an important contributor to dementia. However, little is known about its molecular effects on the human brain and how these alterations may contribute to dementia. Here, we investigated these questions using large-scale quantification of over 8300 proteins from 438 post-mortem brains from a discovery and replication cohort. A proteome-wide association study and protein network analysis of cerebral atherosclerosis found 114 proteins and 5 protein co-expression modules associated with cerebral atherosclerosis. Enrichment analysis of these proteins and modules revealed that cerebral atherosclerosis was associated with reductions in synaptic signaling and RNA splicing and increases in oligodendrocyte development and myelination. A subset of these proteins (n=23) and protein modules (n=2) were also associated with Alzheimer’s disease (AD) dementia, implicating a shared mechanism with AD through decreased synaptic signaling and regulation and increased myelination. Notably, neurofilament light (NEFL) and medium (NEFM) chain proteins were among these 23 proteins, and our data suggest they contribute to AD dementia through cerebral atherosclerosis. Together, our findings offer insights into effects of cerebral atherosclerosis on the human brain proteome, and how cerebral atherosclerosis contributes to dementia risk.

---

Alzheimer’s disease (AD) is the most common cause of dementia and its hallmark pathologies include β-amyloid and neurofibrillary tangles^1^. Both β-amyloid and neurofibrillary tangle pathologies have been studied extensively and found to contribute to synaptic loss, mitochondrial dysfunction, alterations in neuronal activity, and neurodegeneration^1–3^. Yet, it is increasingly recognized that multiple pathologies contribute to the clinical diagnosis of AD dementia in addition to its hallmark pathologies^4–8^. Notably, cerebral atherosclerosis is an important contributor to the development of AD dementia^8–11^ and associated with approximately three times higher risk for AD dementia and nine times higher risk for vascular dementia^12–15^. Despite the high prevalence of cerebral atherosclerosis and its contribution to stroke^16^, AD^8–11^, and vascular dementia^12–15^, we have limited insights into its molecular effects on the human brain, and how these contribute to dementia.

Cerebral atherosclerosis is the accumulation of cholesterol-laden plaques in the walls of large arteries leading to effects ranging from minor wall thickening to significant luminal stenosis and reduced blood flow to the brain^17,18^. Cerebral atherosclerosis is a common condition with a prevalence that increases with age and with the presence of vascular risk factors such as hyperlipidemia, hypertension, diabetes, or smoking^17–19^. For example, it is present in 7% of asymptomatic middle-aged individuals and 82% of individuals over 80 years old ^14,20,21^. Furthermore, cerebral atherosclerosis and its consequences, including lower brain perfusion and metabolism, are among the earliest changes associated with development of AD dementia^8–11^.

To investigate molecular effects of cerebral atherosclerosis on the human brain and how these contribute to dementia, we generated large-scale multiplex brain proteomic quantification of 8356 proteins in 391 subjects from the Religious Order Study and Rush Memory and Aging Project (ROS/MAP; discovery dataset) and 6533 proteins in 47 subjects from the Baltimore Longitudinal Study of Aging (BLSA; replication dataset). These individuals underwent detailed neuropathological assessment of nine age-associated brain pathologies - β-amyloid, neurofibrillary tangles, cerebral atherosclerosis, gross infarcts, microinfarcts, cerebral amyloid angiopathy (CAA), TDP-43, Lewy body, and hippocampal sclerosis – allowing for the identification of the proteomic signature of cerebral atherosclerosis independently of the eight other measured pathologies.

We found that cerebral atherosclerosis was associated with alterations in 114 proteins and 5 protein co-expression modules independently of the eight other measured brain pathologies. Gene set enrichment analysis of these altered proteins and modules revealed that cerebral atherosclerosis was associated with decreased synaptic signaling and plasticity, less RNA splicing and processing, and more oligodendrocyte differentiation, development, and re-myelination.

To investigate how cerebral atherosclerosis contributes to AD dementia, we examined the overlap between proteins and protein modules that contribute to AD dementia and cerebral atherosclerosis independently of the other eight measured brain pathologies. We found that cerebral atherosclerosis likely contributes to AD through reductions in synaptic signaling, plasticity, and regulation but increases in abundances of proteins involved in myelination, possibly in response to axonal injury.

Notably, we found that abundances of neurofilament light (NEFL) and medium (NEFM) chain proteins were associated with cerebral atherosclerosis but not with any of the other eight measured pathologies. Specifically, increased levels of brain NEFL and NEFM proteins were associated with greater severity of cerebral atherosclerosis independently of the other eight measured brain pathologies in both the discovery and replication datasets. Furthermore, increased levels of NEFL and NEFM proteins were associated with AD dementia in both the discovery and replication datasets, suggesting that NEFL and NEFM contribute to AD dementia through cerebral atherosclerosis.

Together, our novel findings provide insights into the direct molecular effects of cerebral atherosclerosis on the human brain and how these can increase the risk of dementia. Furthermore, our findings provide promising protein targets for future studies to identify novel surrogate brain biomarkers for early detection of cerebral atherosclerosis and AD dementia.

## METHODS

### Study design and participants for the discovery dataset (ROS/MAP cohorts)

A total of 391 participants were from two longitudinal clinical-pathologic cohort studies of aging and Alzheimer’s disease - Religious Orders Study and Rush Memory and Aging Project (ROS/MAP) cohorts (Supplementary Table 1a) ^22^. ROS is comprised of older Catholic priests, nuns, and monks throughout the USA. MAP recruited older lay persons from the greater Chicago area. Both studies involve detailed annual cognitive and clinical evaluations and brain autopsy. Participants provided informed consent, signed an Anatomic Gift Act, and a repository consent to allow their data and biospecimens to be repurposed. The studies were approved by an Institutional Review Board of Rush University Medical Center.

### Cognitive diagnosis and brain pathologies in the discovery dataset (ROS/MAP cohorts)

Final clinical diagnosis of cognitive status, including no cognitive impairment, mild cognitive impairment (MCI), Alzheimer’s disease (AD) dementia, or dementia due to other causes, was determined at the time of death by a neurologist with expertise in dementia using all available clinical data but blinded to postmortem data. The diagnosis of dementia was based on the recommendation of the National Institute of Neurological and Communicative Disorders and Stroke and the AD and Related Disorders Association^23^. MCI was based on accepted criteria. Case conferences including one or more neurologists and a neuropsychologist were used for consensus as necessary.

Cerebral atherosclerosis was pathologically assessed by visual inspection of the vessels in the Circle of Willis including vertebral, basilar, posterior cerebral, middle cerebral, and anterior cerebral arteries and their proximal branches^15^. Severity of atherosclerosis was graded with a semiquantitative scale on the basis of the extent of involvement of each artery and the number of arteries involved. Scores ranged from 0 to 3 and were treated as a semiquantitative variable in our analyses. A score of zero means no significant atherosclerosis observed. A score of 1 (mild) indicates that small amounts of luminal narrowing were observed in up to several arteries without significant occlusion. A score of 2 (moderate) means that luminal narrowing occurred in up to half of all visualized major arteries with less than 50% occlusion of any single vessel. Lastly, a score of 3 (severe) indicates that luminal narrowing occurred in more than half of all visualized arteries, and/or more than 75% occlusions of one or more vessels^15^.

Presence of chronic gross infarcts was determined by neuropathologic evaluation blinded to clinical data and reviewed by a board-certified neuropathologist in nine regions (midfrontal, middle temporal, entorhinal, hippocampal, inferior parietal, anterior cingulate cortices, anterior basal ganglia, thalamus, and midbrain) and treated as a dichotomous variable of present or absent ^24^. Presence of microinfarcts in nine brain regions was determined by neuropathological evaluation blinded to clinical data and reviewed by board-certified neuropathologist and treated as a dichotomous variable ^25^.

Neurofibrillary tangles and β-amyloid were identified by molecularly specific immunohistochemistry and quantified by stereology and image analysis, respectively, in eight brain regions – hippocampus, entorhinal cortex, midfrontal cortex, inferior temporal, angular gyrus, calcarine cortex, anterior cingulate cortex, and superior frontal cortex ^26^. Tangle density was determined using systematic sampling and was the average of the tangle densities in eight regions. β-amyloid score represents the percent area of the cortex occupied by β-amyloid and is calculated by taking the mean of β-amyloid scores in 8 brain regions. To create approximately normal distribution of tangles and β-amyloid, we took the square root of tangle density and β-amyloid score, respectively, in our analyses.

Lewy body pathology was assessed using antibodies to α-synuclein in nine regions and were scored based on four stages of distribution of α-synuclein in the brain including 0 (not present), 1 (nigral-predominant), 2 (limbic-type) and 3 (neocortical-type), and treated as present or absent in our analyses ^27^. Hippocampal sclerosis was identified as severe neuronal loss and gliosis in the hippocampus or subiculum using haematoxylin and eosin stain and treated as present or absent in our analyses ^28^. TDP-43 cytoplasmic inclusions were assessed in six regions using antibodies to phosphorylated TDP-43 ^29^. Inclusions in each region were rated on a six-point scale and the mean of the regional scores was created. TDP-43 scores were dichotomized into absent (i.e. mean score of 0) or present (mean score >0) in our analyses. Cerebral amyloid angiopathy (CAA) was assessed in the midfrontal, midtemporal, angular, and calcarine cortices using immunostain for β-amyloid ^30^. Scores were averaged across these 4 regions and treated as a continuous measure.

### Proteomic profiling for the discovery dataset (ROS/MAP cohorts)

The dorsolateral prefrontal cortex was chosen for examination because neuropathologic changes associated with AD are relatively late features of the disease in that region of the brain. We posit that this will enable detection of earlier changes in the brain proteome with disease. Post-mortem tissues from the dorsolateral prefrontal cortex (Brodmann area 9) was cortically dissected. Procedures for tissue homogenization were performed essentially as described previously ^31^. Approximately 100 mg (wet tissue weight) of brain tissue was homogenize in 8 M urea lysis buffer (8 M urea, 10 mM Tris, 100 mM NaHPO4, pH 8.5) with HALT protease and phosphatase inhibitor cocktail (ThermoFisher) using a Bullet Blender (NextAdvance). Each Rino sample tube (NextAdvance) was supplemented with ~100 μL of stainless-steel beads (0.9 to 2.0 mm blend, NextAdvance) and 500 μL of lysis buffer. Tissues were added immediately after excision and samples were then placed into the bullet blender (in 4 °C cold room). The samples were homogenized for 2 full 5 min cycles and the lysates were transferred to new Eppendorf Lobind tubes. Each sample was then sonicated for 3 cycles consisting of 5 s of active sonication at 30% amplitude followed by 15 s on ice. Samples were then centrifuged for 5 min at 15,000 g and the supernatant was transferred to a new tube. Protein concentration was determined by bicinchoninic acid (BCA) assay (Pierce). For protein digestion, 100 μg of each sample was aliquoted and volumes normalized with additional lysis buffer. An equal amount of protein from each sample was aliquoted and digested in parallel to serve as the global pooled internal standard (GIS) in each TMT batch described below. Samples were reduced with 1 mM dithiothreitol (DTT) at room temperature for 30 min, followed by 5 mM iodoacetamide (IAA) alkylation in the dark for another 30 min. Lysyl endopeptidase (Wako) at 1:100 (w/w) was added and digestion continued overnight. Samples were then 7-fold diluted with 50 mM ammonium bicarbonate (AmBic). Trypsin (Promega) was then added at 1:50 (w/w) and digestion was carried out for another 16 h. The peptide solutions were acidified to a final concentration of 1% (vol/vol) formic acid (FA) and 0.1% (vol/vol) triflouroacetic acid (TFA) and desalted with a 30 mg HLB column (Oasis). Each HLB column was rinsed with 1 mL of methanol, washed with 1 mL 50% (vol/vol) acetonitrile, and equilibrated with 2×1 mL 0.1% (vol/vol) TFA. The samples were then loaded and each column was washed with 2×1 mL 0.1% (vol/vol) TFA. Elution was performed with 2 rounds of 0.5 mL of 50% (vol/vol) acetonitrile.

### Isobaric tandem mass tag (TMT) peptide labeling in the discovery dataset (ROS/MAP cohorts)

Prior to TMT labeling, samples were randomized by co-variates (age, sex, PMI, cognitive diagnosis, and pathologies) into 50 total batches (8 cases per batch). Peptides from each individual case (N=400) and the GIS pooled standard (N=100) were labeled using the TMT 10-plex kit (ThermoFisher). In each batch, TMT channels 126 and 131 were used to label GIS standards, while the 8 middle TMT channels were reserved for individual samples following randomization. Labeling was performed as described previously ^31,32^. Briefly, each sample (100 μg of peptides each) was re-suspended in 100 μL of 100 mM TEAB buffer. The TMT labeling reagents were equilibrated to room temperature and 256 μL anhydrous acetonitrile was added to each reagent channel and softly vortexed for 5 min and 41 μL of the corresponding TMT channels were transferred to peptide suspensions. The samples were then incubated for 1 h at room temperature. The reaction was quenched with 8 μl of 5% (vol/vol) hydroxylamine. All 10 channels were then combined and dried by SpeedVac to approximately 150 μL and diluted with 1 mL of 0.1% (vol/vol) TFA and then acidified to a final concentration of 1% (vol/vol) FA and 0.1% (vol/vol) TFA. Sep-Pak desalting was performed with a 200 mg tC18 Sep-Pak column (Waters). Each Sep-Pak column was activated with 3 mL of methanol, washed with 3 mL of 50% (vol/vol) acetonitrile, and equilibrated with 2×3 mL of 0.1% TFA. The samples were then loaded and each column was washed with 2×3 mL 0.1% (vol/vol) TFA, followed by 2 mL of 1% (vol/vol) FA. Elution was performed with 2 rounds of 1.5 mL of 50% (vol/vol) acetonitrile. The elution was dried to completeness.

### High-pH off-line fractionation in the discovery dataset (ROS/MAP cohorts)

High pH fractionation was performed as previously described^33^ with slight modification. Dried samples were re-suspended in high pH loading buffer (0.07% vol/vol NH4OH; 0.045% vol/vol formic acid, 2% vol/vol acetonitrile) and loaded onto an Agilent ZORBAX 300Extend-C18 column (2.1mm × 150 mm with 3.5 µm beads). An Agilent 1100 HPLC system was used to carry out the fractionation. Solvent A consisted of 0.0175% (vol/vol) NH4OH; 0.01125% (vol/vol) formic acid; 2% (vol/vol) acetonitrile and solvent B consisted of 0.0175% (vol/vol) NH4OH; 0.01125% (vol/vol) formic acid; 90% (vol/vol) acetonitrile. The sample elution was performed by a 58.6 min gradient with a flow rate of 0.4 mL/min. The gradient goes 100% solvent A for 2 min, from 0% to 12% solvent B in 6 min, from 12% to 40 % over 28 min, from 40% to 44% in 4 min, from 44% to 60% in 5 min, and then kept 60% solvent B for 13.6 min. A total of 96 individual fractions were collected across the gradient and subsequently pooled into 24 fractions and dried to completeness by speedvac.

### Mass spectrometry analysis in the discovery dataset (ROS/MAP cohorts)

All fractions were resuspended in equal volume of loading buffer (0.1% formic acid, 0.03% trifluoroacetic acid, 1% acetonitrile) and analyzed by liquid chromatography coupled to mass spectrometry essentially as described^34^ with slight modifications. Peptide eluents were separated on a self-packed C18 (1.9 um Dr. Maisch, Germany) fused silica column (25 cm × 75 μM internal diameter; New Objective, Woburn, MA) by a Dionex UltiMate 3000 RSLCnano liquid chromatography system (ThermoFisher Scientific) and monitored on an Orbitrap Fusion mass spectrometer (ThermoFisher Scientific). Sample elution was performed over a 180-min gradient with flow rate at 225 nL/min. The gradient goes from 3% to 7% buffer B in 5 mins, from 7% to 30% over 140 mins, from 30% to 60% in 5 mins, 60% to 99% in 2 mins, kept at 99% for 8 min and back to 1% for additional 20 min to equilibrate the column. The mass spectrometer was set to acquire in data dependent mode using the top speed workflow with a cycle time of 3 seconds. Each cycle consisted of 1 full scan followed by as many MS/MS (MS2) scans that could fit within the time window. The full scan (MS1) was performed with an m/z range of 350-1500 at 120,000 resolution (at 200 m/z) with AGC (automatic gain control) set at 4×10^5^ and maximum injection time 50 ms. The most intense ions were selected for higher energy collision-induced dissociation (HCD) at 38% collision energy with an isolation of 0.7 m/z, a resolution of 30,000 and AGC setting of 5×10^4^ and a maximum injection time of 100 ms. Five of the 50 TMT batches were run on the Orbitrap Fusion mass spectrometer using the SPS-MS3 method as previously described^31^.

### Database Searches and protein quantification in the discovery dataset (ROS/MAP cohorts)

All raw files were analyzed using the Proteome Discoverer suite (version 2.3 ThermoFisher Scientific). MS2 spectra were searched against the canonical UniProtKB Human proteome database (Downloaded February 2019 with 20,338 total sequences). The Sequest HT search engine was used and parameters were specified as: fully tryptic specificity, maximum of two missed cleavages, minimum peptide length of 6, fixed modifications for TMT tags on lysine residues and peptide N-termini (+229.162932 Da) and carbamidomethylation of cysteine residues (+57.02146 Da), variable modifications for oxidation of methionine residues (+15.99492 Da) and deamidation of asparagine and glutamine (+0.984 Da), precursor mass tolerance of 20 ppm, and a fragment mass tolerance of 0.05 Da for MS2 spectra collected in the Orbitrap (0.5 Da for the MS2 from the SPS-MS3 batches). Percolator was used to filter peptide spectral matches (PSM) and peptides to a false discovery rate (FDR) of less than 1%. Following spectral assignment, peptides were assembled into proteins and were further filtered based on the combined probabilities of their constituent peptides to a final FDR of 1%. In cases of redundancy, shared peptides were assigned to the protein sequence in adherence with the principles of parsimony. Reporter ions were quantified from MS2 or MS3 scans using an integration tolerance of 20 ppm with the most confident centroid setting.

### Quality control of proteomic profiles in the discovery dataset (ROS/MAP cohorts)

For each batch, the global internal standards were used to check for proteins outside of the 95% confidence interval and set to missing. Proteins with missing values in more than 50% of the 400 subjects were excluded. Each protein abundance was then scaled by a sample-specific total protein abundance to remove effects of protein loading differences, and then log_2_ transformed. Outlier samples were identified and removed through iterative principal component analysis (PCA). In each iteration, samples more than four standard deviations from the mean of either the first or second principal component were removed, and principal components were recalculated for the next iteration. A total of 9 subjects were removed after three rounds of PCA. After quality control, 391 subjects having 8356 proteins were included in our analyses. The normalized abundance of these 8356 proteins was the residual of the linear regression to remove effects of protein batch, MS2 versus MS3 reporter quantitation mode, sex, age at death, postmortem interval, and study (ROS vs. MAP). The normalized abundance of these proteins was used in our analyses.

### Estimate of brain cell type composition in the discovery dataset (ROS/MAP cohorts)

To account for the heterogeneity of cell types in brain tissues in our analyses, we estimated the proportions of neuron, astrocyte, oligodendrocyte, and microglia using CIBERSORT pipeline and Sharma’s cell type-specific proteins as the reference ^35,36^. We modified the CIBERSORT pipeline by taking the absolute values of the negative regression coefficients instead of setting them to zero. This minor modification produced estimates of cell types similar to those obtained using the original CIBERSORT pipeline and RNA sequencing data. We included the proportion of each of these four cell types as covariates in our proteomic analyses.

### Weighted protein co-expression network analysis in the discovery dataset (ROS/MAP cohorts)

Weighted protein co-expression network analysis was performed using Weighted Gene Co-expression Network Analysis (WGCNA) R package on normalized protein abundance in which effects of batch, sex, age at death, PMI, and study were removed ^37^. We applied the following parameters: soft-thresholding power of 12, “bicor” correlation type, “signed” network type, “signed” Topological Overlap Matrix, a minimum module size of 30, and p-value ratio threshold for reassigning proteins between modules of 0.05, a low propensity to merge modules with “mergeCutHeight” of 0.07, and a high sensitivity to cluster splitting (deepSplit=4). Among the 8356 proteins, 62% (5199 proteins) were assigned to 31 modules, with the largest module having 518 proteins and the smallest module having 46 proteins.

We defined hub proteins as those in the top 20% with regard to high connectivity. To determine the hub proteins for each module, the signed eigenprotein based connectivity, which is the correlation of the protein with its corresponding module eigenprotein, was calculated using the signedKME function from WGCNA package with the parameter corFnc “bicor” ^38^.

### Enrichment analysis for pathways and gene ontology in ROS/MAP

Gene set enrichment analysis of protein co-expression modules was performed using GO-Elite with the background set to all 8356 proteins quantified in the ROS/MAP cohorts ^39^. Protein list per module was subjected to Fisher exact test and Z-test in the Python command line version of GO-Elite (v1.2.5) against the current annotation database for Gene Ontology Biological Process, Molecular Function, Cellular Component, Wiki Pathways, KEGG terms, and Sharma cell types downloaded in January 2019 ^36^. Similarly, gene set enrichment analysis of the cerebral atherosclerosis-associated proteins was performed using the GO-Elite software as described above.

### Clinical and pathologic characterization of the Baltimore Longitudinal Study of Aging cohort (replication dataset)

The replication cohort was 47 subjects from the BLSA with brain proteomic data (Supplementary Table 1b). BLSA is a prospective cohort study of aging in community dwelling individuals ^40^. It continuously recruits healthy volunteers aged 20 or older and follows them for life regardless of changes in health or functional status. The BLSA study was approved by the Institutional Review Board and the National Institute on Aging. Cerebral atherosclerosis was assessed by visual examination of the vessels in the Circle of Willis as described previously ^41^. Areas of atherosclerosis were identified and then sectioned to identify the degree of stenosis by visual inspection and rated on a scale of minimal (<10%), mild (10% <mild<50%), moderate (50%≤moderate<90%), or severe (≥90%). Cerebral atherosclerosis was treated as a semi-quantitative variable in our analyses. Estimates of β-amyloid positive plaques and neurofibrillary tangles were described in detail before by O’Brien and colleagues ^42^. Briefly, silver staining was performed to assess severity of neuritic plaques using CERAD (score ranges from 0 to 3) and of neurofibrillary tangles using Braak (score ranges from 1 to 6), with higher scores indicating higher level of severity. Additionally, cerebral amyloid angiopathy was assessed and rated as mild (rare staining), moderate (scattered, incomplete staining), and severe (circumferential, diffuse staining) and treated as a semi-quantitative variable. Gross infarcts were assessed and quantified as absent versus present anywhere in the brain. Likewise, microinfarcts were assessed and determined as absent versus present in any locations of the brain.

### Proteomic profiling, quality control, and normalization in the BLSA replication cohort

Proteins were profiled from the dorsolateral prefrontal cortex using an analysis pipeline consisting of isobaric tandem mass tag (TMT) mass spectrometry and offline prefractionation as described in detail previously ^32^. TMT labeling was performed following manufacturer’s protocol. MS/MS spectra were searched against a Uniprot human database and Proteome Discoverer 2.1 (ThermoFisher Scientific) as described in detail previously ^32^. Global internal standard (GIS) mixture was checked for extreme outliers, and values outside of the range of log_2_(0.01) to log_2_(100) were excluded from analysis ^32^. Proteins with more than four unquantifiable batches (out of a total of 8 batches) due to 0 or NA value for the GIS channel were excluded from consideration. Proteins with missing value in more than 50% of the samples were excluded. After quality control, 6533 proteins were detected ^32^. Effects of sex, age at death, batch, and postmortem interval were regressed out using non-parametric bootstrap regression ^32^. The normalized proteomic abundance was used for our analyses.

### Statistical analysis

A proteome-wide association study (PWAS) of clinical AD dementia (no cognitive impairment versus MCI/AD dementia), neurofibrillary tangles, β-amyloid, and cerebral atherosclerosis was performed, individually, using linear regression and the normalized protein abundance, adjusting for sex, age at death, proportions of neuron, astrocyte, oligodendrocyte, microglia, and other covariates as specified. Of note, we adjusted for sex, age at death, and cell type composition in all the PWASs. Meta-analysis of the findings from the discovery (ROS/MAP) and replication (BLSA) datasets was performed with METAL using effect size estimates and standard errors^43^.

Associations between protein co-expression modules and clinical AD dementia, neurofibrillary tangles, β-amyloid, and cerebral atherosclerosis, individually, were examined using linear regression, adjusting for sex, age at death, estimated proportions of neuron, astrocyte, oligodendrocyte, microglia, and other covariates as specified. Again, we adjusted for sex, age at death, and cell type composition in all the protein co-expression module analyses. For all analyses, multiple testing adjustment was addressed with Benjamini-Hochberg (BH) false discovery rate (FDR) ^44^.

## RESULTS

### Characteristics of the study cohorts

The discovery dataset consisted of 391 ROS/MAP participants with proteomic data and brain pathological assessment. All were Caucasian, 70% were female, and 99.7% had normal cognition at the time of entering the study. Participants were followed for an average of 8.7 years and median of 9 years (Supplementary Table 1a). The final cognitive diagnosis included normal cognition for 41%, MCI for 26%, AD dementia for 31%, and other dementia for 2% (Supplementary Table 1a). Nine pathologies were assessed post-mortem: β-amyloid, neurofibrillary tangles, cerebral atherosclerosis, gross infarcts, microinfarcts, cerebral amyloid angiopathy (CAA), TDP-43, Lewy body, and hippocampal sclerosis. At the time of death, 32% had gross infarcts, 33% had microinfarcts, and the median cerebral atherosclerosis was mild (i.e. score of 1), with a range of 0 to 3 (Supplementary Table 1a). There was a positive correlation between cerebral atherosclerosis and gross infarcts (r = 0.22; p = 1.60× 10^−5^; Supplementary Figure 1) and a small inverse correlation between cerebral atherosclerosis and β-amyloid (r = −0.13; p = 0.01; Supplementary Figure 1). However, there was no significant correlation between cerebral atherosclerosis and microinfarcts (p = 0.84) or tangles (p = 0.79; Supplementary Figure 1).

The replication dataset consisted of 47 BLSA participants with proteomic data and neuropathological assessment. All were Caucasian, 36% were female, 42% had a diagnosis of AD dementia, and 57% had normal cognition at death (Supplementary table 1b). Six pathologies were assessed post-mortem in these participants: β-amyloid, neurofibrillary tangles, cerebral atherosclerosis, gross infarcts, microinfarcts, and CAA (Supplementary table 1b). Participants had a mean and median cerebral atherosclerosis level of 1.2 (slightly more than mild) and 1 (mild), respectively, with severity range of 0 to 3 (Supplementary table 1b).

### Proteomic analysis of AD dementia and its hallmark pathologies

A total of 8356 proteins remained after quality control. A PWAS of cognitive diagnosis (i.e. no cognitive impairment versus MCI or AD dementia) in the discovery dataset (ROS/MAP cohorts) identified 856 proteins differentially expressed at proteome-wide adjusted p<0.05 (Figure 1A, Supplementary table 2). Among these proteins, those with higher abundance in MCI/AD dementia were enriched for pentose phosphate metabolism, while those with lower abundance were enriched for mitochondrial protein translation (Supplementary figure 2). A PWAS of β-amyloid and neurofibrillary tangles, each adjusting for the 8 other measured pathologies, found 29 and 244 proteins differentially expressed at proteome-wide adjusted p<0.05, respectively (Figure 1B, Supplementary Table 3, and Figure 1C, Supplementary Table 4). The 29 proteins associated with β-amyloid were enriched for proteoglycan metabolic processes (Supplementary figure 3). For neurofibrillary tangles-associated proteins, higher-abundance proteins in greater tangle burden were enriched for regulation of lipid translocation and apoptosis while lower-abundance proteins were enriched for mitochondrial translation (Supplementary figure 4).

**Figure 1:**
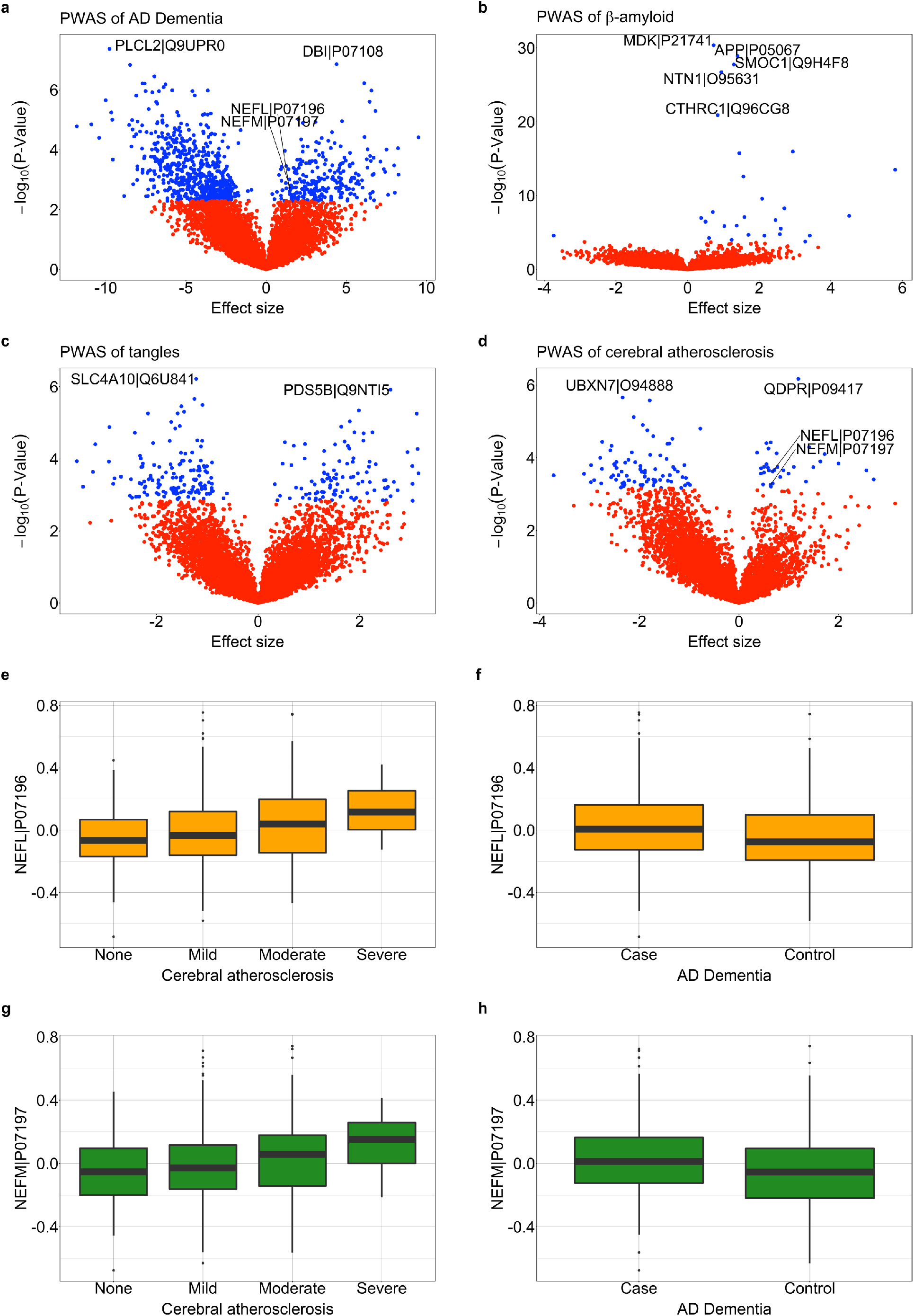
This figure summarizes the proteome-wide association studies of **a)** clinical AD dementia, **b)** β-amyloid, **c)** neurofibrillary tangles, **d)** cerebral atherosclerosis in the discovery dataset (ROS/MAP cohorts; N=375). Each neuropathology examined (**b-d**) was adjusted for the other 8 measured pathologies. Proteins with proteome-wide significant associations were shown in blue, with non-significant associations were shown in red, and the top proteins were labeled. Box plots of **e**) NEFL protein abundance versus cerebral atherosclerosis, **f)** NEFL protein abundance versus AD dementia, **g)** NEFM protein abundance versus cerebral atherosclerosis, **h)** NEFM protein abundance versus AD dementia. For each box plot, the box reflects the first and third quartile, the dark horizontal line reflects the median, the vertical lines extending from the boxes show the 1.5× the interquartile range, and points beyond the lines are outliers.

Network analysis using WGCNA identified 31 protein co-expression modules in the discovery dataset (ROS/MAP cohorts; Figure 2A), and 14 modules were associated with cognitive diagnosis (Figure 2C; Table 1A). Gene set enrichment analysis of these 14 modules revealed that MCI/AD dementia were associated with i) decreased synaptic signaling, transmission, regulation, and plasticity, ii) decreased energy generation and protein synthesis in the mitochondria, iii) increased inflammation, increased myelination, and iv) increased neuronal RNA splicing (Figure 2C; Table 1A). Neurofibrillary tangles were associated with 9 protein modules independently of the 8 other measured pathologies. Furthermore, these 9 modules were also associated with cognitive diagnosis, suggesting that tangles contribute to AD dementia via these modules through i) decreased protein synthesis and energy generation in the mitochondria; ii) increased peptide and neuronal signaling, iii) decreased synaptic transmission; iv) increased inflammation; and v) increased neuronal RNA splicing (Figure 2C; Table 1A). We found that β-amyloid was not associated with any particular protein co-expression modules independently of the 8 other measured pathologies.

**Figure 2:**
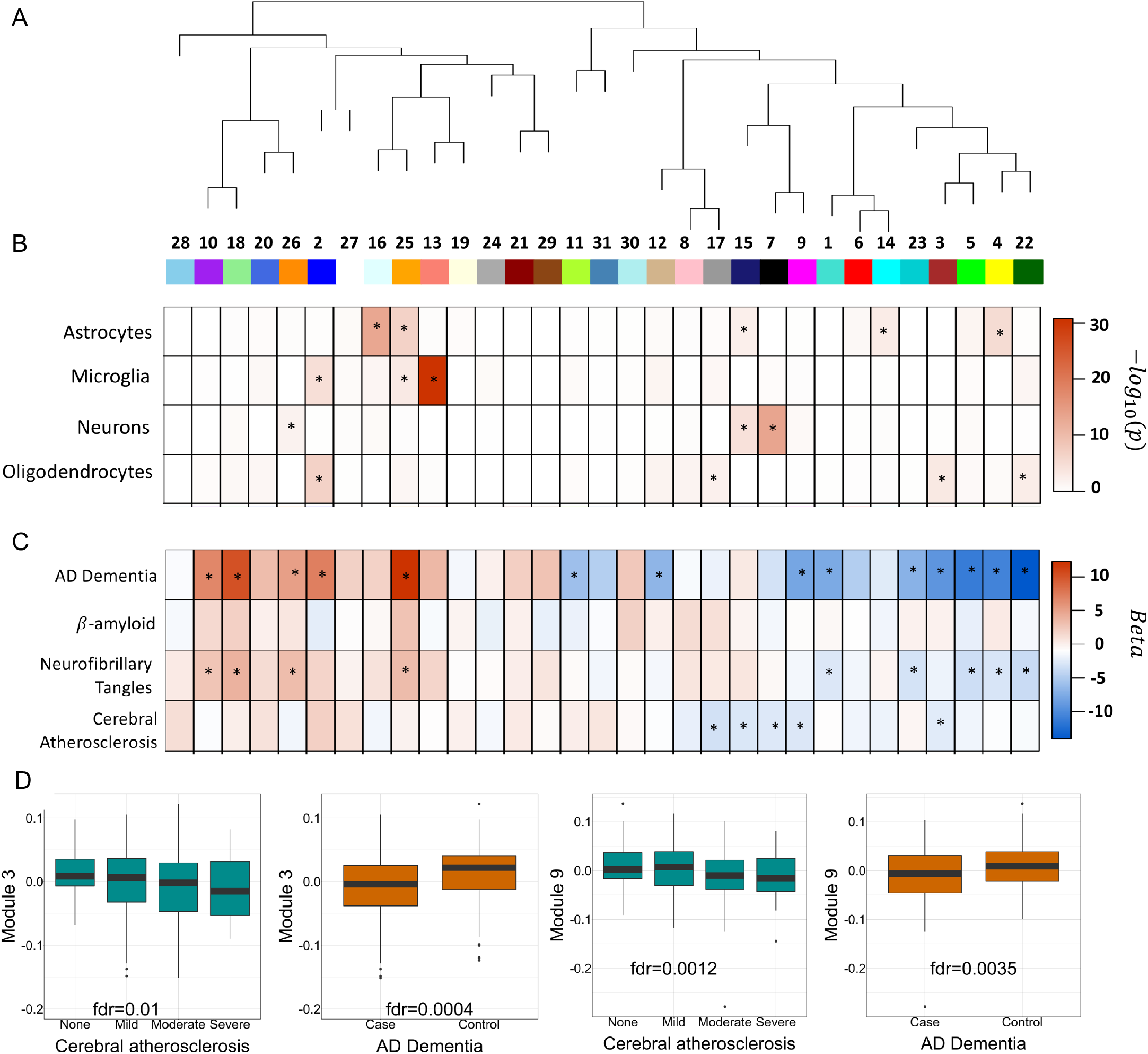
This figure summarizes the protein co-expression modules and their associations with clinical AD dementia, β-amyloid, neurofibrillary tangles, and cerebral atherosclerosis in the discovery dataset (ROS/MAP cohorts). **a)** WGCNA-derived dendrogram for 31 protein co-expression modules. **b)** Brain cell type enrichment was given for each module. **c)** Associations with the outcomes of clinical AD dementia, β-amyloid, neurofibrillary tangles, and cerebral atherosclerosis at adjusted p<0.05 were indicated for each module. No module was associated with β-amyloid after adjusting for all the other 8 measured pathologies at adjusted p<0.05. Nine modules were associated with tangles after adjusting for the 8 other measured pathologies at adjusted p<0.05. Five modules were associated with cerebral atherosclerosis after adjusting for the 8 other measured pathologies at adjusted p<0.05. Beta values on the right-hand bar are the coefficients from the regression model with a dementia feature as the outcome, modules as the predictor, adjusting for the covariates mentioned above. **d)** Box plots of module eigen proteins versus cerebral atherosclerosis and AD dementia to highlight consistent changes in direction of the modules 3 and 9 for both outcomes.

**Table 1:**
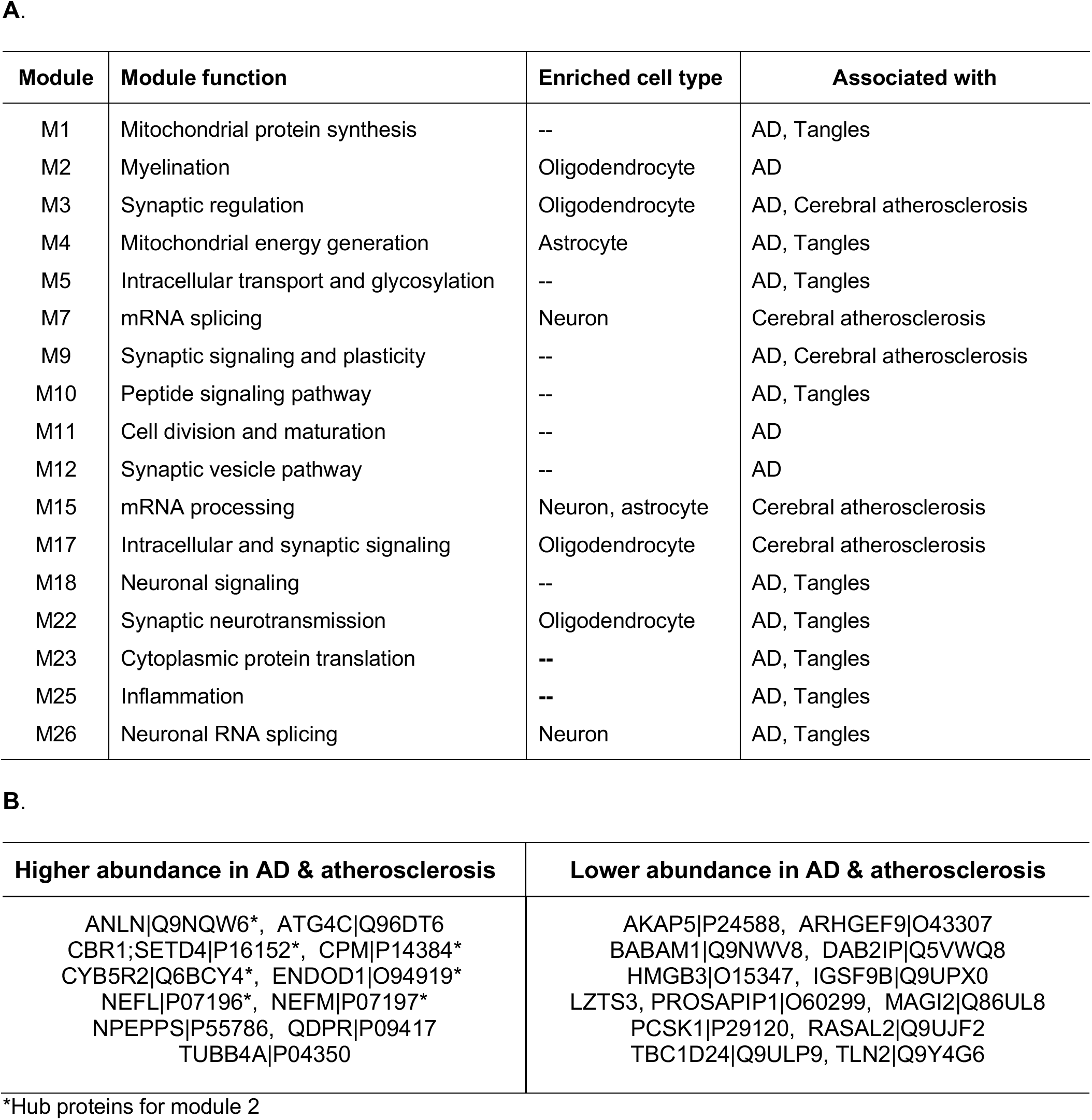
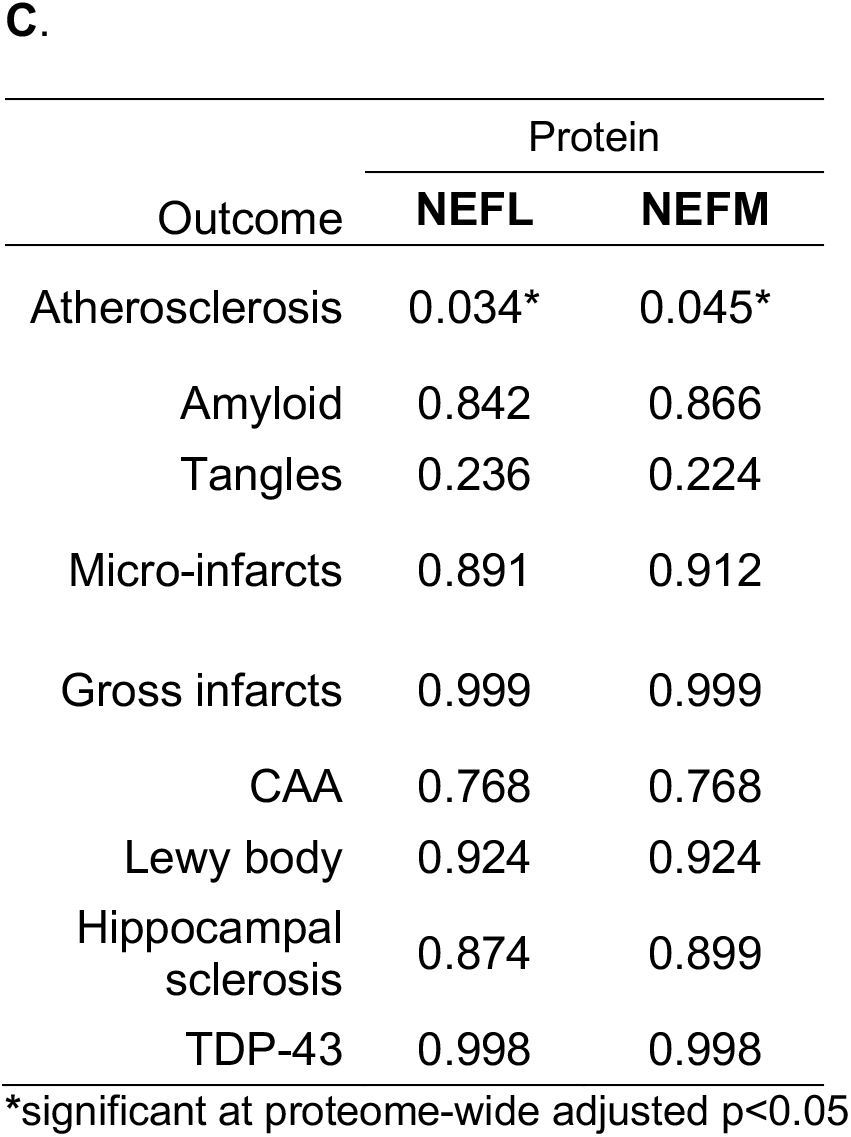
This display item summarizes the proteins and protein modules that were associated with AD dementia, tangles, or cerebral atherosclerosis. **a)** Protein co-expression modules associated with AD dementia, neurofibrillary tangles, or cerebral atherosclerosis in the discovery dataset (ROS/MAP cohorts). All associations presented were significant at adjusted p<0.05 and results of both pathologies were adjusted for the other 8 measured pathologies. **b)** Proteins (n=23) associated with cerebral atherosclerosis independently of the 8 other measured pathologies and with AD dementia at proteome-wide adjusted p<0.05 in the discovery dataset. **c)** Associations between NEFL and NEFM protein levels and each of the 8 pathologies at proteome-wide adjusted p-values in the discovery dataset.

### Brain proteomic signature of cerebral atherosclerosis

The molecular effects of cerebral atherosclerosis on the human brain are not well understood despite its prevalence and contribution to stroke^16^ and dementia^8–15^. To identify proteins altered in cerebral atherosclerosis, a PWAS of cerebral atherosclerosis was performed adjusting simultaneously for the presence of 8 other measured pathologies in the discovery dataset (ROS/MAP cohorts). Cerebral atherosclerosis was defined as a semi-quantitative variable (none/minimal, mild, moderate, or severe) based on assessment of the vessels in the Circle of Willis. There were 114 proteins differentially expressed in cerebral atherosclerosis at a proteome-wide adjusted p<0.05 (Figure 1D, Supplementary table 5). Proteins with positive effect sizes (e.g., QDPR, NEFL, NEMF) had higher abundance and those with negative beta values (e.g., UBXN7) had lower abundance in more severe cerebral atherosclerosis (Figure 1D). Gene set enrichment analysis of the 32 proteins with higher abundance in more severe cerebral atherosclerosis revealed enrichment for oligodendrocyte cell-type specific markers and for oligodendrocyte differentiation, development, and re-myelination (Figure 3). The 82 lower-abundance proteins in greater cerebral atherosclerosis were enriched for RNA splicing and mRNA processing (Figure 3).

**Figure 3:**
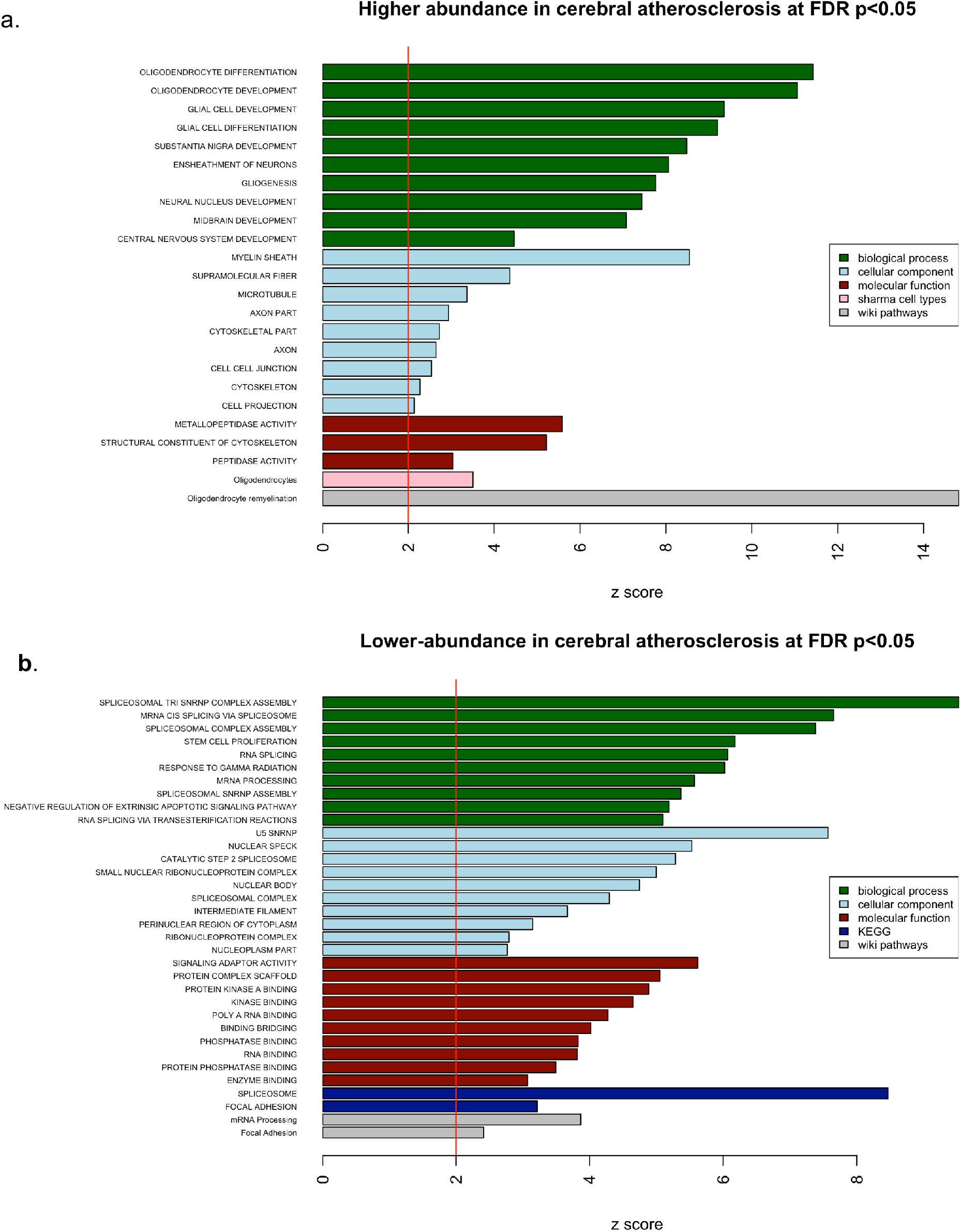
This figure presents the gene set enrichment analysis of the 114 proteins associated with cerebral atherosclerosis independently of the 8 other measured pathologies at proteome-wide adjusted p<0.05 found in the discovery dataset (ROS/MAP cohorts). **a)** Proteins with higher-abundance (n=32) in cerebral atherosclerosis. **b)** Proteins with lower-abundance (n=82) in cerebral atherosclerosis.

Three sensitivity analyses were performed to determine the specificity of our findings for cerebral atherosclerosis. These analyses examined whether gross infarcts or microinfarcts (i.e., the consequence of cerebral atherosclerosis) or vascular risk factors (i.e., predisposing factors for atherosclerosis) were driving the changes in our initial analysis, or whether cerebral atherosclerosis itself was exerting these effects on the brain proteome. In the first analysis, we performed a PWAS of gross infarcts adjusting for the 8 other measured pathologies, which did not yield any significant proteins. Next, we performed a PWAS of cerebral atherosclerosis that did not adjust for gross infarcts but adjusted for all of the other 7 measured pathologies and found more proteins associated with cerebral atherosclerosis (n=237 proteins; Supplementary table 6). However, these proteins were enriched for the same biological and molecular processes as those found in the fully adjusted model for the PWAS of cerebral atherosclerosis (Supplementary Figure 5). In the second sensitivity analysis, we performed a PWAS of microinfarcts adjusting for the 8 other measured pathologies but did not find any significant proteins. Next, we performed a PWAS of cerebral atherosclerosis that did not adjust for microinfarcts but adjusted for all of the other 7 measured pathologies and found more proteins associated with cerebral atherosclerosis (n=142 proteins; Supplementary table 7). Likewise, these 142 proteins were enriched for the same biological and molecular processes as those found in the fully adjusted model (Supplementary Figures 6). In the third sensitivity analysis, we performed a PWAS of vascular risk factors that did not yield any significant proteins. The burden of vascular risk factor here was the sum of counts for lifetime diabetes, hypertension, and smoking, which can contribute to cerebral atherosclerosis ^17,45^. Next, we performed a PWAS of cerebral atherosclerosis adjusting for all 8 measured pathologies and the burden of vascular risk factors. We found essentially the same proteins (106/114) associated with cerebral atherosclerosis after adjusting for vascular risk factors (Supplementary Table 8). Taken together, these sensitivity analyses indicate that the proteomic changes associated with cerebral atherosclerosis in our primary analysis were specific to cerebral atherosclerosis and independent of gross infarcts, microinfarcts, and vascular risk factors.

Among the protein co-expression modules in ROS/MAP cohorts, we found that five modules were associated with cerebral atherosclerosis independently of the other 8 measured pathologies at adjusted p<0.05 (Figure 2C, Table 1A). All five protein co-expression modules had lower abundance in more severe cerebral atherosclerosis (modules 3, 7, 9, 15, and 17; Figure 2C). Among these five protein modules, three modules were enriched for synaptic signaling, regulation, and plasticity (modules 3, 9, and 17), and two modules for mRNA splicing and mRNA processing (modules 7 and 15; Table 1A). Modules 3 and 17 were also enriched for oligodendrocyte cell-type specific markers, modules 7 and 15 for neuron-specific markers, and module 15 also for astrocyte-specific markers (Figure 2B, Table 1A). Taken together, the PWAS and protein co-expression network analysis highlight that cerebral atherosclerosis was associated with i) more oligodendrocyte differentiation, development, and remyelination, ii) less RNA splicing and mRNA processing in neurons and astrocytes, and iii) decreased synaptic signaling, regulation, and plasticity, independently of the 8 other measured pathologies.

To replicate the cerebral atherosclerosis findings in the discovery dataset (ROS/MAP cohorts), we examined these 114 cerebral atherosclerosis-associated proteins in a replication dataset (BLSA cohort). We chose to perform a targeted replication and not a full meta-analysis for all proteins due to the sample size difference between the datasets (391 in the discovery versus 47 in the replication datasets) and technical differences in proteomic profiles due to a selection of proteolytic enzymes (i.e. 8356 proteins in the discovery versus 6533 proteins in the replication), leading to 51% of proteins of the discovery dataset measured in both datasets (4254 of 8356 proteins). Of the 114 cerebral atherosclerosis-associated proteins in the discovery dataset, 79 proteins were measured in the replication dataset. Of these 79 proteins, 42% (33 / 79) were replicated at adjusted p<0.05 (eight proteins had higher abundance and 26 had lower abundance, Supplementary Table 9). All analyses were adjusted for β-amyloid, tangles, gross infarcts, microinfarcts, CAA, sex, age at death, and cell type composition in the same fashion as was done for the discovery dataset. Thus, we were able to replicate 42% of the cerebral atherosclerosis-associated proteins from the discovery dataset (ROS/MAP cohorts) in the replication dataset (BLSA cohort).

### Molecular processes linking cerebral atherosclerosis to elevated odds for dementia

We found that 23 proteins were at the intersection of proteins associated with both cognitive diagnosis and cerebral atherosclerosis independently of the other 8 measured pathologies at proteome-wide adjusted p<0.05 (Table 1B; Supplementary table 10). All 23 proteins consistently had either higher abundance in both MCI/AD and more severe cerebral atherosclerosis, or lower abundance in both, supporting the notion that cerebral atherosclerosis has a detrimental effect on cognition (Supplementary table 10). Of these 23 proteins, 48% (11 of 23) were hub proteins of protein co-expression modules: seven hub proteins for module 2 and one hub protein each for modules 3, 5, 9, and 18 (Table 1B, Supplementary table 10). All the seven hub proteins of module 2 had higher abundance in cerebral atherosclerosis (ANLN, CBR1, CPM, CYB5R2, ENDOD1, NEFL, NEFM; Table 1B, Supplementary Table 10). Module 2 was enriched for myelination and oligodendrocytes cell-type specific markers (Table 1A). Hence, it can be inferred that an aspect of cerebral atherosclerosis’ contribution to clinical AD dementia is through increased abundance of the hub proteins involved in myelination.

Likewise, from the protein network analyses, we identified two modules (3 and 9) that were associated with both cognitive diagnosis and cerebral atherosclerosis independently of the other 8 measured pathologies. Both modules had reduced abundance in more severe cerebral atherosclerosis and in MCI/AD (Figure 2D). Module 3 was enriched for synaptic regulation and oligodendrocyte specific markers, and module 9 for synaptic signaling and plasticity (Table 1A). Together, converging findings from the PWAS and protein network analyses suggest that cerebral atherosclerosis likely contributes to AD dementia though its effects linked to reductions in synaptic signaling, regulation, and plasticity, and increases in abundance of the hub proteins involved in myelination. One possible model explaining these findings is that atherosclerosis causes axonal dysfunction and increase in myelination is in response to axonal injury. Axonal injury is notably supported by the association between greater abundance of neurofilament light (NEFL) and neurofilament medium (NEFM) chain proteins and higher levels cerebral atherosclerosis because NEFL and NEFM are well-described markers of axonal injury^46,47^.

### NEFL and NEFM proteins are associated with cerebral atherosclerosis and AD dementia

Among the 23 proteins associated with both cognitive diagnosis and cerebral atherosclerosis independently of the other 8 measured pathologies, NEFL and NEFM were noteworthy since prior work has shown that their CSF levels associated with AD (Table 1B) ^48–50^. In light of these findings, we examined whether NEFL and NEFM were associated with cerebral atherosclerosis specifically or with other measured pathologies as well. Therefore, we performed a PWAS of each of the remaining pathologies (i.e. gross infarcts, microinfarcts, CAA, TDP-43, Lewy Body, and hippocampal sclerosis) adjusting for the other 8 measured pathologies in the discovery dataset. For each PWAS, we adjusted for all of the remaining 8 measured pathologies to find the proteins specifically associated with the pathology of interest. Strikingly, NEFL and NEFM were not associated with β-amyloid, tangles, microinfarcts, gross infarcts, CAA, Lewy body, hippocampal sclerosis, or TDP-43 (Table 1C). NEFL and NEFM were only associated with cerebral atherosclerosis (NEFL p=0.00024, proteome-wide adjusted p=0.034; NEFM p=0.00056, proteome-wide adjusted p=0.045; Figure 1D,E,G; Table 1C). Associations between NEFL and NEFM proteins and cerebral atherosclerosis were replicated in the replication dataset (BLSA cohort; Supplementary Table 9). These findings underscore the specific associations between NEFL and NEFM protein abundances and cerebral atherosclerosis in both the discovery and replication datasets. Likewise, the associations between NEFL and NEFM protein abundances and cognitive diagnosis in the discovery dataset (NEFL p=0.0025, proteome-wide adjusted p=0.032, OR=1.47; NEFM p=0.0019, proteome-wide adjusted p=0.028, OR=1.51; Figure 1A,F,H) was confirmed in the replication dataset (BLSA dataset: NEFL p=0.016, OR = 5.2; NEFM p=0.023, OR = 4.6). Notably, the associations between NEFL and NEFM proteins with cognitive diagnosis were no longer significant when we adjusted for cerebral atherosclerosis in the regression model, suggesting that cerebral atherosclerosis mediates the association between NEFL, NEFM and AD dementia (NEFL p=0.0076, proteome-wide adjusted p = 0.068; NEFM p=0.0057, proteome-wide adjusted p = 0.058). Taken together, these findings underscore that NEFL and NEFM likely contribute to AD dementia through cerebral atherosclerosis.

## DISCUSSION

We investigated the direct molecular effects of cerebral atherosclerosis on the human brain and how these alterations increase the risk for AD dementia using the deep proteomes (n=8356 proteins) in a large sample size (438 individual post-mortem brains). We found that cerebral atherosclerosis was associated with altered abundance of more than 100 proteins and five protein co-expression modules after adjusting for microinfarcts, gross infarcts, β-amyloid, neurofibrillary tangles, CAA, TDP-43, Lewy body, and hippocampal sclerosis. Gene set enrichment analysis of these proteins and modules revealed that cerebral atherosclerosis was associated with decreased RNA splicing and mRNA processing, lower synaptic signaling and plasticity, and more oligodendrocyte differentiation, development, and remyelination, possibly due to myelin sheath damage. Additionally, we showed that these alterations in cerebral atherosclerosis were not likely the result of either i) risk factors for cerebral atherosclerosis (i.e. hypertension, diabetes, and smoking) or ii) the consequences of cerebral atherosclerosis (i.e., gross or microinfarcts in the brain). Of the proteins associated with cerebral atherosclerosis in the ROS/MAP discovery cohort, nearly half were replicated in the BLSA replication cohort.

An important aspect of cerebral atherosclerosis that we aimed to understand is its contribution to AD dementia. We found 23 proteins and two protein co-expression modules that contribute to both cerebral atherosclerosis and AD dementia independently of β-amyloid, tangles, gross infarcts, microinfarcts, and other pathologies. Thus, the effect of cerebral atherosclerosis on AD dementia is likely through reduced synaptic signaling, reduced plasticity, and injury to the myelin sheath. Our findings are consistent with prior studies in rodent models showing that cerebral hypoperfusion, a direct consequence of cerebral atherosclerosis, was associated with white matter inflammation and oxidative stress, which in turn lead to damage to the myelin sheath and demyelination ^51^. We did not find a statistical interaction between cerebral atherosclerosis and β-amyloid, or between cerebral atherosclerosis and tangles with regard to AD dementia, suggesting that cerebral atherosclerosis mainly contributes to AD dementia independently of the two AD hallmark pathologies; in other words, its effects are primarily additive to those of β-amyloid and neurofibrillary tangles.

Strikingly, protein abundances of NEFL and NEFM in the brain were specifically associated with cerebral atherosclerosis and AD dementia in the discovery dataset (ROS/MAP cohorts) and confirmed in the replication dataset (BLSA cohort). These findings indicate that NEFL and NEFM likely contribute to AD dementia through cerebral atherosclerosis. In support of these findings, another longitudinal cohort (i.e. Banner Sun Health Systems) with brain proteomic data also showed that higher protein abundances of NEFL and NEFM in the brain were associated with faster cognitive decline in advanced age, consistent with their associations with MCI/AD in this study^52^. Also consistent with our findings that NEFL and NEFM act through cerebral atherosclerosis on AD dementia is the lack of association between CSF NEFL and CSF β-amyloid despite the association between CSF NEFL and AD dementia in prior studies in CSF ^49,53^. In addition, we found that NEFL and NEFM were both hub proteins of protein co-expression module 2, which was enriched for myelination, the process in which oligodendrocytes myelinate the axons by wrapping them with myelin sheath. This is consistent because NEFL and NEFM are primarily expressed in myelinated axons ^46,49^. Our findings provide insights into why NEFL is a promising biomarker for AD dementia and its progression as shown by studies of CSF and plasma NEFL in AD ^49,50,54,55^.

From a translational viewpoint, intra-brain vascular dysregulation has been suggested to be the strongest and earliest brain pathology associated with AD dementia development, followed by Aβ deposition, dysregulation of glucose metabolism, and grey matter atrophy in a recent multifactorial analysis of 7,700 brain images and plasma and CSF biomarkers from the Alzheimer’s Disease Neuroimaging Initiative (ADNI) ^10^. Likewise, several studies suggested that reduction or dysregulation of cerebral blood flow occurs before cognitive decline, brain atrophy, and β-amyloid accumulation ^9^. These studies underscore the important and early contribution of cerebral atherosclerosis to the pathogenesis of AD dementia. Hence, surrogate brain biomarkers for cerebral atherosclerosis might be promising and more sensitive for early detection of AD dementia. We provided here 23 brain proteins specifically associated with cerebral atherosclerosis independently of the other 8 measured pathologies and with cognitive diagnosis (Table 1B) for future studies to investigate whether CSF or plasma levels of these proteins can serve as early biomarkers for cerebral atherosclerosis or AD dementia.

In addition to examining the effects of cerebral atherosclerosis on the brain proteome and its potential effects on AD dementia, this work also investigated the proteomic alterations found in AD dementia and 8 other measured pathologies in a large dataset of individuals with deep proteomic profiling (n=383, n=8356 proteins). We found that MCI/AD was associated with decreased synaptic activities and plasticity, decreased mitochondrial protein synthesis and energy generation, increased inflammation, increased myelination, and increased neuronal RNA splicing. These findings are consistent with prior proteomic studies in AD and cognitive decline ^32,52,56^. In addition, we found that neurofibrillary tangles contribute to AD through decreased mitochondrial protein synthesis and energy production, decreased synaptic transmission, increased inflammation, and increased neuronal RNA splicing independently of β-amyloid and the other 7 pathologies.

Interpretation of our findings should take into consideration its limitations. First, this is a postmortem brain study, therefore, it is a cross-sectional study and no causal relationship can be established. Second, the proteomes of the replication dataset (BLSA cohort) had fewer proteins than those of the discovery dataset (ROS/MAP cohorts), reflecting the advances in proteomic technology but making the replication study not as ideal. Our study has several strengths. It is the first human brain PWAS of cerebral atherosclerosis to the best of our knowledge. Second, the human brain proteomes were deep (over 8300 proteins) and the sample size large (N=391). Using isobaric tandem mass tag mass spectrometry, we detected 8356 proteins after quality control, which is more than two times the number of proteins detected from label-free quantification^52^. Third, the discovery and replication datasets have detailed neuropathological assessments, which enabled us to rigorously control for the effects of other co-morbid pathologies and improve the specificity of the protein alterations in cerebral atherosclerosis. Fourth, the discovery and replication design reduces the risk of false positive. Lastly, the discovery dataset (ROS/MAP cohorts) is a community-based cohort and not a convenience sample recruited from cognitive clinics, making the findings more generalizable.

In conclusion, this is the first human brain PWAS to characterize molecular effects of cerebral atherosclerosis on the human brain, and the first study to elucidate molecular underpinnings for contribution of cerebral atherosclerosis to AD dementia. Importantly, we show that protein levels of NEFL and NEFM in the brain were specifically associated with cerebral atherosclerosis and not the other 8 measured pathologies and likely contribute to AD through cerebral atherosclerosis.

## ACKNOWLEDGEMENTS

We thank the participants of the ROS, MAP, and BLSA studies for their time and participation. Support was provided by R01 AG056533, R01 AG053960, the Accelerating Medicine Partnership for AD (U01 046152; U01 AG046161; U01 AG061356; U01 AG061357), the Emory Alzheimer’s Disease Research Center (P50 AG025688), and the NINDS Emory Neuroscience Core (P30 NS055077), R01 AG017917 (D.A.B.), R01 AG015819 (D.A.B.), RC2 AG036547 (D.A.B.), P30 AG10161 (D.A.B.), R01 AG042210 (J.A.S), and in part by the intramural program of the National Institute on Aging (NIA).

A.P.W. is also supported by U01 MH115484 and I01 BX003853. T.S.W. is also supported by NIH grants RF1 AG057470, U01 AG061357, P50 AG025688, R56 AG062256, R56 AG060757, and R01 AG056533. N.T.S. is also supported by an Alzheimer’s Association, Alzheimer’s Research UK, The Michael J. Fox Foundation for Parkinson’s Research, the Weston Brain Institute Biomarkers Across Neurodegenerative Diseases Grant (11060), R01 AG061800, and R01 AG057911.

